# Quantitative multiplex immunohistochemistry reveals inter-patient lymphovascular and immune heterogeneity in primary cutaneous melanoma

**DOI:** 10.1101/2021.10.08.463675

**Authors:** Julia Femel, Jamie L. Booth, Tina G. Asnaashari, Sancy A. Leachman, Takahiro Tsujikawa, Kevin P. White, Young H. Chang, Amanda W. Lund

## Abstract

**Purpose:** Quantitative, multiplexed imaging is revealing complex spatial relationships between phenotypically diverse tumor infiltrating leukocyte populations and their prognostic implications. The underlying mechanisms and tissue structures that determine leukocyte distribution within and around tumor nests, however, remain poorly understood. While presumed players in metastatic dissemination, new preclinical data demonstrates that blood and lymphatic vessels (lymphovasculature) also dictate leukocyte trafficking within tumor microenvironments and thereby impact anti-tumor immunity. Here we interrogate these relationships in primary human cutaneous melanoma.

**Experimental Design:** We established a quantitative, multiplexed imaging platform to simultaneously detect immune infiltrates and tumor-associated vessels in formalin-fixed paraffin embedded patient samples. We performed a discovery, retrospective analysis of 28 treatment-naïve, primary cutaneous melanomas.

**Results:** Here we find that the lymphvasculature and immune infiltrate is heterogenous across patients in treatment naïve, primary melanoma. We categorized five lymphovascular subtypes that differ by functionality and morphology and mapped their localization in and around primary tumors. Interestingly, the localization of specific vessel subtypes, but not overall vessel density, significantly associated with the presence of lymphoid aggregates, regional progression, and intratumoral T cell infiltrates.

**Conclusions:** We describe a quantitative platform to enable simultaneous lymphovascular and immune infiltrate analysis and map their spatial relationships in primary melanoma. Our data indicate that tumor-associated vessels exist in different states and that their localization may determine potential for tumor cell exit (metastasis) or leukocyte trafficking (immune response).

This platform will support future efforts to map tumor-associated lymphovascular evolution across stage, assess its prognostic value, and stratify patients for adjuvant therapy.

**TRANSLATIONAL RELEVANCE:** This report describes a quantitative, image-based method to investigate the relationship between the tumor-associated lymphovasculature and immune landscape in treatment naïve, primary human melanoma. The research shows that melanoma-associated blood and lymphatic vessels display context-dependent phenotypes that associate both with the risk of regional progression and immune infiltration. These findings indicate that stromal/vascular heterogeneity may underlie regional differences in immunogenicity and thus present opportunities for future biomarker development and therapeutic intervention.

## INTRODUCTION

Immuno-oncology is radically changing the clinical landscape for melanoma patients and placing new emphasis on personalizing treatment to optimize response. Biomarkers capable of stratifying risk and response are necessary to facilitate precision in clinical implementation in the adjuvant and neoadjuvant setting (1). Histological evaluation of tumor-infiltrating lymphocytes (TIL) within and around tumors has emerged as one such biomarker yielding insight into immunologic responsiveness (2,3) and providing prognostic power in solid tumors (4). From this growing body of work, it is now clear that not just the number but also the location of TIL is critical in their prognostic value (4,5). The success of these approaches has highlighted a need to understand the underlying mechanisms that determine TIL localization, which may provide both new prognostic biomarkers and orthogonal targets for combination immunotherapy.

Multiplexed immunohistochemistry (mIHC) allows for an unprecedented understanding of the *in situ* biology of multiple cell types and tissue features while preserving structural and spatial relationships (6–8). While much effort has been made to deeply profile the location and functional phenotype of infiltrating leukocytes (6–8), inter- and intra-lesional heterogeneity may indicate that intratumoral immune complexity in part depends on stromal tissue biology. We have yet, however, to leverage mIHC approaches to investigate the functional plasticity of non-hematopoietic, stromal cells and therefore lack an understanding for how underlying tissue features collaborate to determine regional immune surveillance and tumor progression.

The tumor-associated lymphovasculature, both hematogenous and lymphogenous, plays critical roles in regulating tumor progression by providing routes for distant and regional metastasis (9,10). Emerging preclinical work, however, expands the view of both blood and lymphatic vessels from passive routes for metastasis (distant and regional respectively) and indicates that vessel functionality impacts immune surveillance (11–13) and immune escape (14–16). Blood vessels can be activated to enhance leukocyte adhesion and tissue infiltration (11,14,17) and lymphangiogenesis may support enhanced antigen presentation in LNs that contributes to local and distal tumor control (12,13,15,18,19). This work generated the hypothesis that the tumor-associated vasculature might determine response and could be therapeutically targeted to enhance immunotherapy. These preclinical observations, however, have yet to be fully validated in human tissues and must still be reconciled with the metastatic effects of tumor-associated angio- and lymphangiogenesis. Therefore, translating into the clinic depends upon methods to capture the lymphovascular plasticity and its relationship to immune infiltration and tumor progression in clinical samples.

To begin to address this gap in knowledge, we present a platform to simultaneously integrate a spatial understanding of tumor-associated inflammation and the lymphovasculature in formalin-fixed paraffin embedded (FFPE) tissue sections. We explore the hypothesis that vessel phenotype shapes early host immune responses to malignancies and thus may provide unique insight into the risk for lesion development and progression. Using a discovery set of primary cutaneous human melanomas with known LN involvement, we quantify vessel composition across and within patients to assess the relationships between vessel heterogeneity, tumor immune context, and regional progression. By multiplexing established endothelial (AQP1, CD34, PDPN, LYVE-1, MECA-79) and pericyte markers (αSMA), we categorize tumor-associated vessels into arterioles, activated capillaries/postcapillary venules, immature neovasculature, high-endothelial venule-like vessels, lymphatic capillaries, and inflamed lymphatic capillaries. We find that the localization and density of specific vessel phenotypes are associated with immune infiltration and progression. Intratumoral activated capillaries are enriched where CD8^+^ T cells infiltrate tumor nests, while dysfunctional or immature peritumoral capillaries are associated with an increased risk for regional metastasis. Our findings support a rationale for deeper phenotypic analysis of tumor-associated vessels within their spatial context, and present a quantitative platform to enable future work evaluating the prognostic potential of endothelial features in predicting immune responsiveness and risk for progression.

## MATERIAL AND METHODS

### Human melanoma samples

Stage I to III human primary melanoma resections were obtained from the OHSU Knight Biolibrary and the OHSU Department of Dermatology research repository (Table I) according to defined inclusion criteria (Supplemental Figure 1). We selected patients with known LN status as a surrogate for poor prognosis. Acquisition and use of human samples were performed in accordance with the Institutional Review Board at OHSU. Thin sections were assessed by a dermatopathologist to confirm presence of melanoma, tumor type (exclusion of tumors other than “superficial spreading” melanoma), tumor thickness, and tissue integrity. Of the samples that made it through quality assessment, 27 had negative and 14 positive sentinel LN biopsies. Due to extensive tissue loss (a function of repeated heat-mediated antigen retrieval and bleaching), an additional 13 samples were excluded from analysis (Supplemental Figure 1).

**Table I.** Clinical characteristics of human cutaneous primary melanoma sample cohort. Sentinel lymph node (sLN), standard deviation (SD).

| **Baseline Characteristics in Cohort at Inclusion** |  |
| --- | --- |
| Number of Patients | 28 |
|  | # (%) |
| Gender |  |
| male | 14 (50) |
| female | 14 (50) |
| Age, years |  |
| range | 34-88 |
| mean (± SD) | 59.8 ± 15.2 |
| sLN status |  |
| neg | 17 (61) |
| pos | 11 (39) |
| Tumor thickness |  |
| range | 1.05-8.4 |
| mean (± SD) | 2.86 ± 1.68 |
| Pathological stage |  |
| Pt2 | 11 (39) |
| Pt3 | 11 (39) |
| Pt4 | 6 (21) |
| Ulceration |  |
| no | 16 (57) |
| yes | 10 (36) |
| no info | 2 (7) |
| Pathological stage (sLN) |  |
| Pn0 | 17 (61) |
| Pn1 | 6 (21) |
| Pn2 | 4 (14) |
| Pn3 | 1 (4) |

### Multiplex Immunohistochemistry (mIHC)

Sequential chromogenic immunohistochemistry was performed as previously described (8,16), using a modified protocol (Supplemental Table I and II). Prior to staining, tissue sections were baked at 60 °C for 30-90 min. Subsequently, sections were bleached with 10% H_2_O_2_ at 65 °C for 10 min. Antigen retrieval was performed in heated Citra Plus solution (HK080-9K, BioGenex) for 20 min. ImmPRESS™ horseradish peroxidase (HRP) polymer reagent (Vector Laboratories) and AEC (Vector Laboratories) were used for detection and visualization. Whole slide imaging was performed on the Aperio ImageScope AT (Leica Biosystems). AEC was removed using ethanol, antibody was stripped in heated Citra solution (except prior to LYVE-1 staining, for which tissues was treated with heated Target Retrieval Solution pH 6.1 (S169984-2, Dako) for 30 min). Hematoxylin staining was performed after the final round. Individual staining patterns were validated by single stains using control sections (e.g. tonsil) and primary antibody removal after each round of antibody staining confirmed.

### Image registration and ROI selection

Serial digitized images were processed using a computational image analysis workflow described previously (7,8). Images were aligned and registered in MATLAB (version R2018b) using the SURF algorithm in the Computer Vision Toolbox (The MathWorks, Inc., Natick, MA) (20). Square-shaped regions of interest (ROIs) with an area of 6.25 mm^2^ were selected for analysis based on presence of PDPN^+^ lymphatic vessels within tissue areas containing stromal tissue and tumor parenchyma. Depending on tissue size, an average of three ROIs/sample were chosen. ROI coordinates were applied to the registered images for each sample (Supplemental Figure 2B). Bleaching of melanin and repeated treatment of slides in heated Citra Plus solution resulted in regional tissue loss. To exclude regions with tissue loss from subsequent analyses, the final hematoxylin staining was used for ROI selection and downstream segmentation.

### Color processing

Image processing was performed using FIJI (“FIJI Is Just ImageJ”) (21). AEC extracted quantification and visualization using a custom macro for color deconvolution (NIH plugin RGB_to_CMYK). The FIJI plugin Color_Deconvolution [H AEC] was used to separate hematoxylin. Color deconvolution was followed by postprocessing steps for signal cleaning and background elimination (22). For visualization color-deconvoluted images were overlaid in Aperio ScanScope (Leica Biosystems) and pseudo-colored.

### Tumor segmentation

Binary tumor segmentation masks were generated to enable analysis within intratumoral and peritumoral regions as previously described (16). In brief, the hematoxylin image was used to define tissue areas using triangle thresholding and S100^+^ area was detected by computing an alternate sequential filter, followed by triangle thresholding. All tumors were S100^+^, variation in staining intensity does not affect segmentation. Single S100^+^ cells, which might include Langerhans cells, dendritic cells, macrophages and neural cells, are not used to define tumor area and instead the analysis is focused on tumor nests. The inverted intratumoral mask was used to define peritumoral tissue area.

### Leukocyte analysis

Analysis of single immune cells was performed with CellProfiler Version 3.5.1 (23). Cell populations were classified based on a hierarchical gating using image cytometry with FCS Express 6 Image Cytometry RUO (De Novo Software, Glendale, CA).

### Vessel segmentation and analysis

Whole-vessel segmentation was performed using Otsu’s method to segment blood and lymphatic vessels based on AQP1, CD34, and PDPN staining. panCK staining was used to exclude PDPN^+^ basal epithelial cells. Single channel images are denoised and locally flattened by using a mathematical morphology operation. The resulting images are used in Otsu’s thresholding, creating a binarized image of vascular structures and size-based filtering applied to resulting structures were used to remove technical artifacts. Subpopulations of vessels were defined using supervised hierarchical gating based on positive pixel coverage and intensity thresholding of the vessel markers (Supplemental Figure 2D; Figure 1B). Thresholds were determined manually and confirmed by visual inspection. Morphological and shape features were extracted.

**Figure 1.**
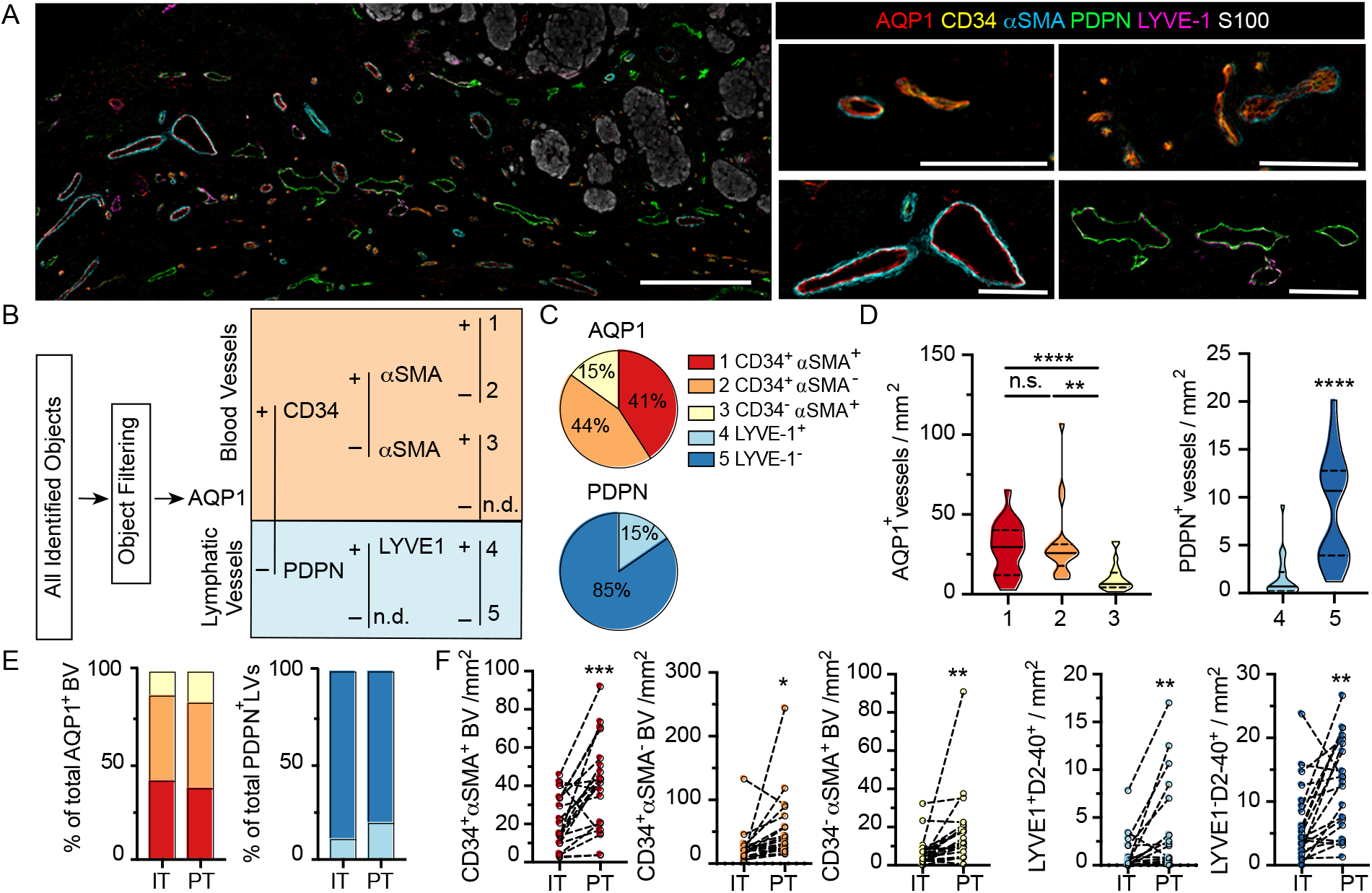
Human primary cutaneous melanomas exhibit a phenotypically heterogenous lymphovasculature. (**A**) Formalin-fixed paraffin embedded thin sections from archived primary cutaneous melanomas were analyzed by multiplexed immunohistochemistry to analyze regional lymphovascular subtypes. Scale bar (left) = 300μm; (right)=100μm. (**B**) Supervised, quantitative vessel analysis was performed to identify vascular objects and evaluate positivity for five lymphovascular markers (AQP1, CD34, αSMA, PDPN, LYVE1) to identify distinct vessel subtypes. (**C**) Relative proportions of each vessel subtype among total blood (AQP1^+^) or lymphatic (PDPN^+^) vessels and (**D**) and densities across the entire tissue area. One-way ANOVA or unpaired students t test. (**E**) Vessel proportions and (**F**) and densities segmented by region (intratumoral, IT; peritumoral, PT). Paired students t-test. *p<0.05, **p<0.01, ***p<0.001.

### Statistics

Statistics were calculated with Graphpad Prism software v8.3 (Graphpad) as indicated in figure legends. Data were tested for normality and tests applied as appropriate (normal, Student’s t test; non-normal, Mann Whitney). One-way ANOVA was used for multiple comparisons. Pearson or Spearman were applied to normal and non-normal data respectively. Data is presented as averages of all ROIs analyzed per sample, as individual ROIs, or as individual vessels as indicated.

## RESULTS

### Human primary cutaneous melanoma exhibits a phenotypically heterogenous lymphovasculature

To directly test the relationship between lymphovascular density and heterogeneity, immune infiltration, and tumor progression, we designed a discovery cohort of treatment naïve, primary human cutaneous melanoma (Table I). Using this cohort we took a multiplexing approach to simultaneously detect one melanoma (S100), one epithelial (pancytokeratin), 4 blood vessel (AQP1, CD34, αSMA, MECA79), 2 lymphatic vessel (PDPN, LYVE-1), and 4 leukocyte (CD45, CD8, CD20, CD68) markers (Supplemental Table I). Lymphovascular markers were selected to first segregate the two major vessel types, blood (aquaporin 1; AQP1) and lymphatic (podoplanin; PDPN), and second to provide insight into their functional phenotype (CD34; alpha smooth muscle actin, αSMA; lymphatic vessel endothelial hyaluronan receptor 1, LYVE-1). While loss of antigenicity of S100 is observed in a subset of metastatic melanomas (1-4%) (24), all patients enrolled in our study showed S100-reactivity. The use of alternative or additional melanoma markers, such as GP100/MART-1, could improve coverage across larger cohorts. Images collected for each individual stain were processed through an established pipeline to segment intratumoral and peritumoral regions, single leukocytes, and vessels (Supplemental Figure 2). The resolution of intratumoral masks was tuned to eliminate the contribution of single, isolated S100^+^ cells. All data was integrated for downstream analyses.

Upon image registration and vessel characteristics quantitation (Supplemental Figure 2), the tremendous heterogeneity of the lymphovasculature within and around melanoma lesions was readily apparent (Figure 1A). Through supervised, hierarchical gating, we identified 5 vessel types, 3 blood (1. AQP1^+^CD34^+^ αSMA^+^; 2. AQP1^+^CD34^+^ αSMA^-^ 3. AQP1^+^CD34^-^ αSMA^+^) and 2 lymphatic (4. PDPN^+^LYVE-1^+^ 5. PDPN^+^LYVE-1^-^) (Figure 1B), which we classify as (1) immature, neovasculature, (2) activated capillaries/postcapillary venules, (3) arterioles, (4) lymphatic capillaries, and (5) inflamed lymphatic capillaries (Supplemental Table I and Supplemental Figure 3A). Vessel classification is supported by single cell RNA sequencing of endothelial cells from human non-small cell lung cancer and normal adjacent tissue (25). Analysis of this publicly available data confirmed broad expression of AQP1 across all blood endothelial cells but not lymphatic endothelial cells (LEC), with enrichment for CD34 on activated capillaries and postcapillary venules (Supplemental Figure 3B and C). While PDPN was evenly expressed by all LECs as expected, normal LECs retained high LYVE-1 expression whereas tumor-associated LECs were largely LYVE-1 negative (Supplemental Figure 3D). Similarly, we observed loss of LYVE-1 expression on PDPN^+^ vessels proximal to melanoma tumor nests. AQP1^+^CD34^-^αSMA^-^ objects were single, S100^+^ and CD45^-^ cells and are likely tumor cells (26). No PDPN^-^LYVE-1^+^panCK^-^ structures were identified. All objects were visually validated to be multicellular, vascular structures.

In addition to their surface phenotype, vessel size and patency may be a surrogate of functionality in tissue sections. Due to a lack of automated, quantitative methods to segment and extract morphological features from multicellular structures, however, studies that explore vessel structure are limited. Therefore, we generated a quantitative image-based vessel segmentation and analysis pipeline to extract morphological features that describe vessel shape, openness, and size (Supplemental Figure 4A). As expected, across vessel subtypes, AQP1^+^ blood vessels were larger and rounder than PDPN^+^ lymphatic vessels, which exhibited a flattened, irregular morphology (Supplemental Figure 4B). Intratumoral AQP1^+^ blood vessels were larger and flatter than peritumoral AQP1^+^ vessels (Supplemental Figure 4C), independent of surface phenotype. This is consistent with expected increases in tumor cellularity, nutrient deprivation-induced intratumoral angiogenesis and elevated intratumoral stresses (27). Intratumoral PDPN^+^ vessels are rare, but when found did not differ morphologically from their peritumoral counterparts (Supplemental Figure 4D).

While AQP1 and PDPN served to classify blood from lymphatic vessels, we additionally used a set of markers to indicate subtype, CD34, αSMA, and LYVE-1. Notably most studies use only one endothelial marker and therefore fail to capture any functional heterogeneity that may exist. CD34 is a transmembrane sialomucin and adhesion molecule that plays a critical role in leukocyte transmigration and thus marks sites of potential leukocyte infiltration into tumors; αSMA is an actin isoform that marks contractile pericytes; and LYVE-1 is a hyaluronan binding receptor implicated in docking and transmigration of dendritic cells and tumor cells. A majority of AQP1^+^ vessels were CD34^+^ (Figure 1C) and these activated capillaries (AQP1^+^CD34^+^ αSMA^+/-^) were found at higher density than AQP1^+^CD34^-^ αSMA^+^ arterioles (Figure 1D). While dense, contractile pericyte investment is expected in arterioles (αSMA^+^), capillaries and post-capillary venules generally exhibit intermittent αSMA^-^ pericyte coverage (Supplemental Figure 3A). Therefore, the abnormal investment of some tumor-associated capillaries with contractile pericytes (immature; AQP1^+^CD34^+^ αSMA^+^) may indicate a dysfunctional phenotype. These vessels exhibited the largest filled area while activated capillaries and postcapillary venules (AQP1^+^CD34^+^ αSMA^-^) appeared flatter (Supplemental Figure 4E). The vast majority of lymphatic vessels detected lacked expression of the canonical marker LYVE-1 (Figure 1C and D). Interestingly, LYVE-1^-^ vessels were smaller and flatter than their LYVE-1^+^ counterparts (Supplemental Figure 4F). Subtype analysis was similar in intratumoral and peritumoral regions (Figure 1E), but all vessel types showed increased density at the tumor periphery relative to intratumoral regions (Figure 1F), indicating that primary melanomas are largely hypovascular relative to their immediately adjacent stroma. These data indicate that tumor-associated vessels exhibit functional and morphological heterogeneity within and around tumor nests in primary melanoma.

### Treatment naïve, primary cutaneous melanoma microenvironments exhibit robust peritumoral inflammation and lymphoid aggregates

In addition to lymphovascular markers we evaluated the density and distribution of key leukocyte populations, including cytotoxic T cells (CD8), B cells (CD20), macrophages (CD68), and other CD45^+^ leukocytes (CD8^-^CD20^-^CD68^-^). Tumor-associated inflammation was largely restricted to the peritumoral region (Figure 2A). The exclusion was most significant for CD20^+^ B cells and other CD45^+^ leukocytes, while both CD8^+^ T cells and CD68^+^ macrophages on average showed a more homogenous distribution between intratumoral and peritumoral regions across the patient samples (Figure 2B).

**Figure 2.**
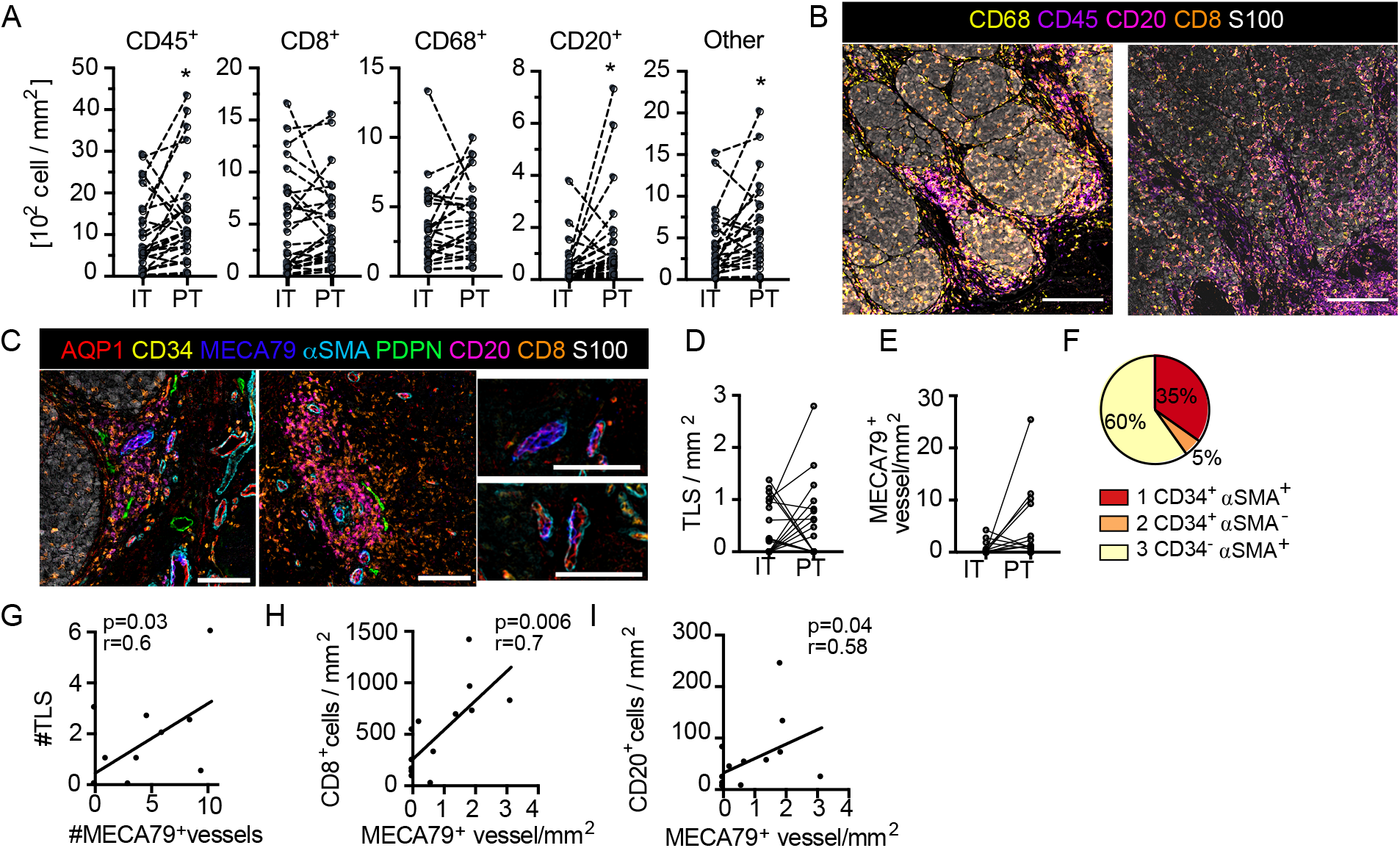
Treatment naïve, primary cutaneous melanoma is characterized by excluded immune infiltrates and lymphoid aggregates. (**A**) Leukocyte density segmented by intratumoral (IT) and peritumoral (PT) regions. Paired student’s t-test. (**B**) Images of immune infiltrates in tumor microenvironments of two representative patients (scale bar = 500µm). (**C**) Representative images of tertiary lymphoid-like structures (TLS) (top; scale bar = 150µm) and associated AQP1^+^MECA79^+^ high-endothelial venule-like vessels (bottom; scale bar = 100µm). (**D**) IT and PT TLS density. (**E**) IT and PT AQP1^+^MECA79^+^ vessel density. (**F**) MECA79^+^ vessels across AQP1^+^ subtypes. Correlation of (**G**) TLS with AQP1^+^MECA79^+^ vessels and (**H**) AQP1^+^MECA79^+^ vessel density with CD8^+^ or (**I**) CD20^+^ cell density. Pearson or Spearman correlations. **p<0.05, ***p<0.001.

One striking feature was the presence of lymphoid clusters at and around the tumor boundary, reminiscent of tertiary lymphoid structures (TLS). We, therefore, added an additional endothelial marker to our panel, MECA79, which binds carbohydrate epitopes on peripheral node addressins, including sulfated ligands for L-selectin (including CD34, GlyCAM1 etc), and promotes T and B lymphocyte adhesion and rolling across high endothelial venules (HEV) in the LN. This specialized vessel state, therefore, defines sites of naïve and memory lymphocyte infiltration into tumors, which has been proposed to be a good prognostic in melanoma (28,29). In our analysis we defined TLS by co-aggregation of CD8^+^ T and CD20^+^ B cells and found that presence of TLS was closely associated with these specialized HEV-like vessels (Figure 2C), as expected. TLS (Figure 2D) and AQP1^+^MECA79^+^ HEV-like vessels were distributed both intratumoral and peritumoral (Figure 2E). Interestingly MECA79 positivity was enriched in vessels with invested αSMA^+^ pericytes (Figure 2F). The number of AQP1^+^MECA79^+^ vessels directly correlated with the number of aggregates observed across patient samples (Figure 2G) and was the best vessel-based positive correlate with overall CD8^+^ T and CD20^+^ B cell density (Figure 2H and I). Interestingly, in all cases, these aggregates were associated with at least one PDPN^+^ lymphatic vessel (Figure 2C). While our sample size is too small to evaluate the correlation of either AQP1^+^MECA79^+^ vessel or aggregates with LN metastasis, it is clear that both LN positive and negative disease can exhibit primary tumor lymphoid aggregates in the treatment naïve setting.

### Regionally metastatic cutaneous primaries are poorly inflamed and exhibit increased immature capillary density

Angiogenesis, lymphangiogenesis, and tumor-associated inflammation are all predicted drivers of early dissemination in solid tumors. To begin to query the relationship between the TME and probability of regional progression, we looked for associations between our image features and presence of LN metastases. While our study is not powered to identify independent prognostic variables, an association of lymphovascular and immune features with progression suggests relevance for future validation in large cohorts. Interestingly, primaries that had not metastasized to regional draining LNs exhibited increased intratumoral leukocyte (CD45^+^) density over regionally metastatic primaries (Figure 3A), however, no individual leukocyte subset (CD20^+^ B cells, CD68^+^ macrophages, or CD8^+^ T cells) independently drove this difference (Figure 3B and C; data not shown). The intratumoral ratio between CD8^+^ and CD68^+^ cells, which is predictive of long-term survival in primary melanoma (6), also failed to associate with regional progression in this cohort (Figure 3D). To further understand the inter-patient heterogeneity in immune context, we looked at the relative densities of each leukocyte type across individual patients (Figure 3E and F). We observed three types of inflammatory environments, patients in subset 1 all had LN negative disease and exhibited robust inflammation with relatively high densities of CD8^+^, CD20^+^, CD68^+^ and other CD45^+^ leukocytes. In contrast a second group was similarly CD8^+^ T cell rich but lacked high densities of other leukocytes. Finally, a poorly inflamed group lacked T and B cells with moderate CD68^+^ macrophage densities. This patient subset exhibited a roughly equal proportion of LN positive and negative disease. These data indicate that a more inflamed TME is negatively associated with regional progression, however, whether a single immune signature could be predictive of LN metastasis remains to be determined in larger cohorts.

**Figure 3.**
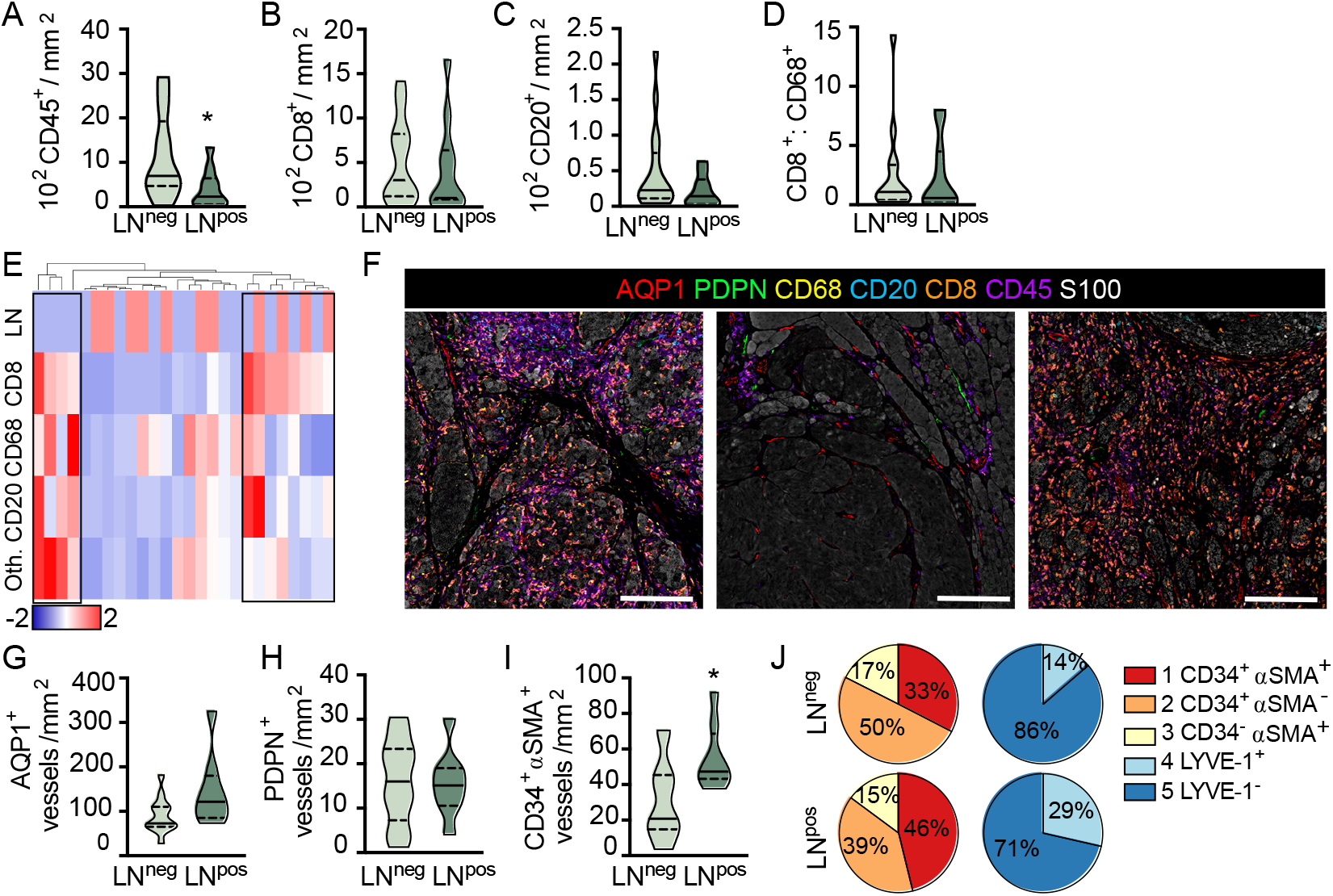
Lymph node metastasis is associated with reduced inflammation and increased immature capillary density. (**A**) Intratumoral CD45^+^ leukocyte, (**B**) CD8^+^ T cell and (**C**) CD20^+^ B cell density as a function of lymph node (LN) status. (**D**) IT CD8^+^:CD68^+^ cell ratio. (**E**) Heatmap representation of leukocyte density (CD8^+^, CD68^+^, CD20^+^, and other [Oth.] CD45^+^) across all ROIs per sample revealing three inflammatory subtypes. Scaled by row, clustered by individual patients (column). Top row, sentinel LN positive (red); LN negative (blue). (**F**) Representative images of inflammatory subtypes. Inflamed (left), uninflamed (middle), and T cell only (right). Scale bar=200μm. (**G**) Peritumoral AQP1^+^ total blood and (**H**) PDPN^+^ total lymphatic vessel density in LN positive and negative primary melanomas. (**I**) Peritumoral density of immature, neovasculature (AQP1^+^CD34^+^ αSMA^+^) in LN negative and positive primary melanomas. (**J**) Vessel subtype heterogeneity as a function of total AQP1^+^ blood or PDPN^+^ lymphatic vessels and LN status. Unpaired student’s t-test, *p<0.05.

We next looked at the lymphovasculature to determine whether vessel density correlated with LN positivity. Neither blood (AQP1^+^) nor lymphatic (PDPN^+^) vessel density was statistically different between LN negative and positive disease, in either peritumoral (Figure 3G and H) or intratumoral regions (Supplemental Figure 5A and B), however, there appeared to be a trend towards increased peritumoral blood vessel density. We therefore asked whether any individual blood vessel subtype explained this trend and found that LN positive disease exhibited a significantly higher density of peritumoral AQP1^+^CD34^+^ αSMA^+^ immature neovasculature (Figure 3I) with no significant change in density observed in either of the other vessel subtypes (data not shown). Interestingly, for each individual patient, those with LN metastases appeared to have an overall higher proportion of immature or dysfunctional AQP1^+^CD34^+^ αSMA^+^ blood capillaries and resting (non-inflamed) PDPN^+^LYVE-1^+^ lymphatic capillaries relative to non-metastatic tumors (Figure 3J and Supplemental Figure 5D), though heterogeneity across patients was clearly evident. In fact, it is interesting to note that even in this treatment naïve setting, significant inter- and intra-patient lymphovascular heterogeneity was revealed, consistent with similar observations in non-small cell lung cancer (25). We found no correlation between CD45^+^ leukocyte infiltrates and AQP1^+^CD34^+^ αSMA^+^ vessel density (Supplemental Figure 5E), perhaps indicating that there will be utility in the combination of these markers to stratify disease. Future work with expanded cohorts will determine the independence of these variables and their prognostic value for predicting both progression and survival.

Finally, primary tumor ulceration is a known adverse prognostic indicator and previously shown to be associated with changes in blood vessel size, patency, and vascular invasion (30). We therefore evaluated our parameters as a function of ulceration (Supplemental Figure 5F). We noted no difference in CD45^+^ leukocyte infiltrate (Supplemental Figure 5G) or any leukocyte subtype (data not shown), nor did we see a difference in AQP1^+^ blood (Supplemental Figure 5H) or PDPN^+^ lymphatic vessel density or heterogeneity (Supplemental Figure 5I). We did, however, consistent with the literature, observe a significant enlargement of AQP1^+^ blood vessels in ulcerated samples with open lumen (Supplemental Figure 5J), specifically within the immature neovasculature and arteriole subsets (Supplemental Figure 5K-M). We saw no evidence of lymphovascular invasion in our cohort. These data taken together indicate that there may be both immune and lymphovascular based determinants of early dissemination. How they collaborate to determine outcomes remains an exciting and open question.

### Intratumoral, activated capillaries are associated with improved regional CD8^+^ T cell infiltration in cutaneous melanoma

Our observation that CD8^+^ T cells were not strictly excluded from intratumoral regions (Figure 2A and B) led us to wonder if vessel density or subtype could provide insight into the phenomenon of T cell exclusion. While physical barriers present at the tumor periphery may sequester or exclude T cells (31), it is also likely that the presence of patent and activated intratumoral blood vessels may be necessary for tumor infiltration; a hypothesis that is used to justify normalizing angiogenesis therapy in combination with immune checkpoint blockade (32). To test this hypothesis, we classified all non-overlapping regions of interest (ROI) as excluded or infiltrated (Figure 4A and B) by determining the ratio of CD8^+^ T cells intratumoral and peritumoral (IT:PT). We analyzed multiple ROIs for most tumors, however, this remains only an estimate of overall tumor behavior given the extensive spatial heterogeneity observed even within a single thin section. To this point, both infiltrated and excluded ROIs were identified in the same lesions, supporting our hypothesis that independent of the systemic lymphocyte repertoire, local stromal biology contributes to tissue infiltration and spatial compartmentalization of immune responses.

**Figure 4.**
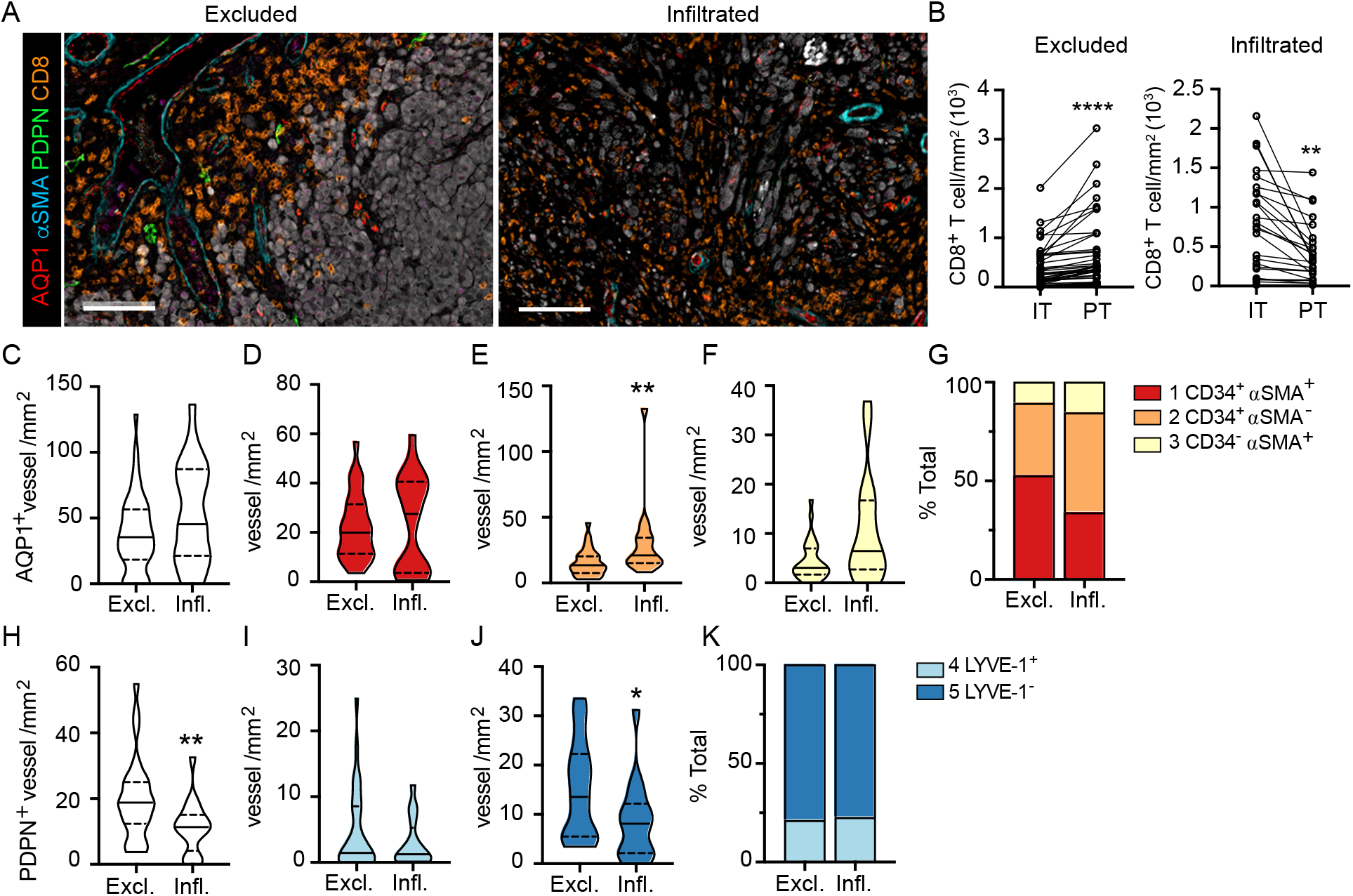
Intratumoral, activated capillaries are associated with improved regional CD8^+^ T cell infiltration in primary melanoma. (**A**) Representative images of CD8^+^ T cell excluded (Excl.) and infiltrated (Infl.) regions of interest (ROI). Scale bar=100μm. (**B**) Intratumoral (IT) and peritumoral (PT) CD8^+^ T cell density in Excl. and Infil. tumor regions. Paired student’s t-test, **p<0.01. (**C**) Total AQP1^+^, (**D**) immature neovasculature (AQP1^+^CD34^+^ αSMA^+^), (**E**) activated capillaries and postcapillary venules (AQP1^+^CD34^+^ αSMA^-)^, and (**F**) arteriole (AQP1^+^CD34^-^ααSMA^+^) density (IT) and (**G**) percent of total. (**H**) Total PDPN^+^, (**I**) lymphatic capillary and (**J**) inflamed lymphatic capillary density (PT) and (**K**) percent of total. Unpaired student’s t-test, *p<0.05.

We found no statistical difference in intratumoral blood vessel (AQP1^+^) density between excluded and infiltrated ROIs (Figure 4C), so we analyzed each subtype and observed a statistically significant increase in the density of activated capillaries (AQP1^+^CD34^+^ αSMA^-^) in infiltrated over excluded ROIs (Figure 4E). We saw no change in the density of other subtypes (Figure 4D and F). This was consistent with an increase in proportion of activated capillaries and a concomitant decrease in proportion of intratumoral AQP1^+^CD34^+^ αSMA^+^ immature neovasculature (Figure 4G), notably the vessel subtype enriched in patients with LN metastasis (Figure 3I). These phenotypic changes were also associated with changes in morphology. Intratumoral vessels in well infiltrated tumors were larger with increased lumen area (Supplemental Figure 6A and B), indicating that a normalized, activated, and patent intratumoral vasculature may support T cell infiltration. Interestingly, when we looked at the presence of peritumoral lymphatic vessels in these same ROIs we observed a significant decrease in overall PDPN^+^ vessel density that was driven by PDPN^+^LYVE-1^-^ inflamed lymphatic capillaries (Figure 4H-K). Interestingly, both types of lymphatic vessels found peritumoral to infiltrated tumors exhibited a flattened morphology with increased eccentricity (Supplemental Figure 6C and D), perhaps indicating reduced mechanical stresses in the stroma (e.g. interstitial fluid pressures or fibrosis) that would drive lymphatic vessel distension (27). These data indicate that vessel morphology and phenotype associate with TIL localization, observations that may have relevance for TME normalizing therapies (32).

## DISCUSSION

Tumor-associated angiogenesis and lymphangiogenesis are both correlated with metastatic potential in human patients and clinical models (9,10). In preclinical models, we and others have further demonstrated that the tumor-associated vasculature has significant implications for immune surveillance, immune infiltration, and immune escape (11,13,15). This has led to the hypothesis that therapeutically targeting lymphovascular function might synergize with immunotherapy and improve patient outcomes (11,12,17,18). Further translation of these ideas, however, requires assays that capture structural, positional, and functional lymphovascular features in patient specimens. To address this need, we have established a multiplexed imaging workflow to capture inter- and intra-patient lymphovascular and immune heterogeneity in FFPE patient tissues. We map lymphovascular phenotype and location, and find that specific vessel subtypes, not overall vessel density (blood or lymphatic), associated with regional progression and local immune infiltration. This work suggests that a deep understanding of the underlying tumor-associated vasculature may provide insight into local immune potential and disease progression that could be leveraged to identify and apply new combinations in the adjuvant setting. While this discovery data set is not powered to evaluate the independent prognostic value of any one feature, the data presented provide the impetus for future work that explores vascular and non-hematopoietic biology in concert with immune infiltrates to generate a more detailed map of the TME and guide clinical care.

Blood vessels provide both a route for distal tumor dissemination and the entry point for tumor-infiltrating leukocytes (9). Lymphatic vessels similarly promote regional spread to satellite cutaneous sites and LNs, while also providing the requisite route for dendritic cell migration, antigen presentation, and immune activation (13,33,34). While these paradoxical roles are beginning to be explored in preclinical models (11,12,14,15,18), there are limited studies that investigate the interrelationship of metastatic potential, immune infiltrate, and the vasculature in human tumor tissue. One of the few studies to do so made the surprising observation that lymphatic vessel density negatively correlated with distal metastasis and positively correlated with CD8^+^ T cell infiltrate, such that a combined lymphatic and T cell score stratified overall survival in colorectal cancer (19). This is consistent with smaller studies that have indicated a regional relationship between lymphatic vessel density and immune infiltrate, suggesting co-regulation, at least in the context of melanoma (16,35). In the dataset presented here, we did not observe a statistically significant relationship between lymphatic or blood vessel density and the overall density of CD8^+^ T cells, but rather, regional phenotypic changes associated with T cell localization. In contrast, a low density of peritumoral lymphatic vessels and high density of intratumoral AQP1^+^CD34^+^ αSMA^-^ activated capillaries and postcapillary venules indicated a TME favorable for intratumoral T cell infiltration. These observations are consistent with preclinical work that demonstrates that vascular phenotype may support or inhibit tumor infiltration (14) and suggests that careful histological analysis of the endothelium in human tumors may be predictive of response to therapy or indicate synergistic therapeutic opportunities (11,17).

Though the dogma states that lymphatic vessel density is associated with increased probability of LN metastasis (36), we did not find a statistical relationship in our cohort. While this may be a consequence of sample size, there is similar discordance across published studies (37), which is consistent with preclinical observations that while lymphangiogenesis may promote LN metastasis (38) it is not required (39). Here we show that there are at least two subsets of lymphatic vessels in cutaneous melanoma, PDPN^+^LYVE-1^+^ and PDPN^+^LYVE-1^-^, where LYVE-1 is reported in some cases to correlate with the extent of LN metastasis and overall poor prognosis (40). This may be consistent with a direct role for LYVE-1 in cell docking and transmigration (41,42), or simply indicate absence of local inflammation (43). We note that poorly inflamed tumors are more regionally metastatic than inflamed tumors. Still, lymphatic vessel number or density does not seem to be a reproducible predictor of regional progression, and it remains to be determined whether further functional markers could reveal additional prognostic value. Interestingly, we instead see an increase in density of immature AQP1^+^ αSMA^+^CD34^+^ vessels in primary tumors that have metastasized to LNs. While these vessels don’t provide the route for LN metastasis, the recruitment of contractile αSMA^+^ pericytes to intratumoral capillaries may indicate abnormal vessel activation and is associated with resistance to anti-angiogenic therapy (44), perhaps indicating a dysfunctional TME.

Ectopic lymphoid aggregates that form in peripheral tissues, termed tertiary lymphoid organs (TLO), often arise in the setting of chronic inflammation, autoimmunity, and cancer (45). TLO are characterized by their similarity to secondary lymphoid organs, with respect to cellular organization and vascular specialization, and can display features of active germinal centers that are hypothesized to be a site of *de novo* antigen presentation and lymphocyte priming. In the setting of solid tumors, these structures are generally a good prognostic, may arise on therapy, and were recently associated with improved response to immune checkpoint blockade in melanoma (28,29). Our data indicates that treatment naïve, cutaneous melanomas already exhibit structures consistent with a TLO-like structure; aggregates of T and B cells associated with MECA79^+^ vasculature. Notably, the structures we identified were highly correlated with overall B and T cell density but were not specifically enriched in either LN positive or negative disease. The prominence of MECA79^+^ vessels in these structures is consistent with the idea that vascular activation is causal for local lymphoid neogenesis (11). We further see that lymphoid aggregates co-localize with PDPN^+^ lymphatic vessels at the tumor border, similar to observations in kidney cancer (46). The association of regional lymphatic vessels with lymphoid aggregates is interestingly consistent with observations that disrupted fluid flows (47,48) and lymphatic-derived chemokines (18,49) may initiate ectopic lymphoid organ development.

Importantly, our work describes a workflow for coincident analysis of lymphovascular and immune components within intact TME and suggests a rationale for deeper phenotypic analysis of vascular heterogeneity and plasticity. This platform could enable future work to compare the primary and metastatic vasculature across solid tumor types. Cutaneous metastases may be a particularly interesting test case given the expected increase in incidence of lymphovascular invasion. Furthermore, what determines early dissemination to lymph nodes rather than systemic dissemination and mortality may be distinct, and future work with annotated distal metastasis and survival data will be critical to begin to tease these two concepts apart.

Finally, the current study is limited by cohort size and is thus ripe for future validation in large, annotated cohorts. Further limitations of our approach include tissue loss over time and variability in staining across batches as a function of both normal variation, antibody lots, and sample heterogeneity. Importantly, bleaching may alter antigen retrieval and staining patterns, and we found CD31 staining to be particularly sensitive to this protocol. Finally, quantitative image analysis harbors inherent biases in ROI selection, tumor segmentation, and data integration that all significantly impact results. While many studies image small regions of interest the advantage of our approach are large ROIs and objective quantitative analysis. Future work should continue to define standard methods and approaches to tissue sampling and quantification.

In conclusion, we build off of preclinical hypotheses to expand the view of the tumor-associated lymphovasculature from a static, homogenous structure to a functional and dynamic component of the immune TME in melanoma. Our work supports the hypothesis that lymphovascular changes within TME may both reflect ongoing immunity and directly contribute to its localization. Incorporating additional markers of endothelial activation into the analysis, a deeper dive into mural cell phenotypes, and image-based functional features will further add value to this platform, providing both rationale for new biological hypotheses and a discovery platform for novel biomarkers.

## Acknowledgements

The authors acknowledge Eric Smith MS, MBA, Pamela Cassidy, PhD, and the OHSU Biolibrary for support in sample identification and acquisition and Ryan S. Lane, PhD for analysis support. This work was funded by the OHSU Knight Cancer Center Support grant NIH P30-CA069533, the Department of Defense Peer Reviewed Cancer Research Program (AWL; W81XWH-15-1-0348), the V Foundation for Cancer Research (AWL: V2015-024), Melanoma Research Alliance (AWL; 403181), and the Cancer Research Institute Clinic and Laboratory Integration Program (CLIP) (AWL; No Number). J.F. acknowledges support from the Swedish Research Council (Vetenskapsrådet, International postdoc grant). YHC acknowledges the support from NIH/NCI U54 CA209988. SAL was supported in part by the OHSU Knight Cancer Institute’s John D. Gray Endowment.

## Author Contributions

J.F. performed experiments, designed the study, and analyzed data. T.G. optimized analysis algorithms. J.L.B. performed experiments. S.A.L. and K.P.W. consulted on sample acquisition, quality control, and data analysis. T.T. established mIHC platform and consulted on technical aspects of the manuscript. Y.H.C. developed vessel segmentation algorithms. A.W.L. obtained funding, designed the study, analyzed data, and wrote the manuscript. All authors reviewed the final manuscript.

## Supplementary Material

**Supplemental Table I.** Antibody product information. LEC: lymphatic endothelial cell.

| Markers | Antibody (host) | Target | Clone | Product # | Manufacturer | Dilution |
| --- | --- | --- | --- | --- | --- | --- |
| *Lymphovascular* | Podoplanin (mouse) | PDPN; mucin-type protein, highly expressed by LECs and necessary for lymphatic development. | D240 | 916601 | BioLegend | 1:100 |
|  | LYVE-1 (rabbit) | Lymphatic vessel endothelial hyaluronan receptor 1; binds both soluble and insoluble hyaluronan, expressed by LECs. | polyclonal | ab36993 | Abcam | 1:100 |
|  | AQP1 (rabbit) | Aquaporin 1; widely expressed water channel showing specificity to blood over lymphatic vessels. | polyclonal | AB2219 | Millipore | 1:10000 |
|  | CD34 (mouse) | Sialomucin, L-selectin ligand. Critical for T lymphocyte rolling and entry into lymph nodes. | QBEnd-10 | MA1-10202 | Pierce (Thermo Fisher) | 1:10000 |
|  | MECA79 (rat) | Binds L-selectin ligands collectively called peripheral lymph node addressins (GlyCAM-1, CD34, Agp200, and endomucin). | IgM | 120801 or 2 | Biolegend | 1:100 |
| | αSMA (rabbit) | Alpha smooth muscle actin; Actin isoform involved in contractile apparatus of smooth muscle. | polyclonal | ab5694 | Abcam | 1:500 |
| *Melanocyte* | S100 (mouse) | Low molecular weight calcium binding proteins expressed by melanocytes. Also expressed in other cells of neural crest origin, macrophages, and dendritic cells. | cocktail | CM089B | Biocare Medical | 1:2000 |
| *Epithelial* | panCytokeratin (mouse) | Keratin proteins found in the cytoskeleton of epithelial tissue, marks epidermis. | [AE1/AE3] | Ab27988 | Abcam | 1:2000 |
| *Immune* | CD45 (mouse) | Pan-leukocyte marker. | HI30 | 14-0459-82 | eBioscience | 1:200 |
|  | CD20 (rabbit) | B-lymphocyte antigen, expressed from late pro-B cells through memory but not plasma cells | SP32 | ab64088 | Abcam | 1:1000 |
|  | CD8 (mouse) | Co-receptor of the T cell receptor, marker of cytotoxic T cells. | C8/144B | MS-457-S1 | Thermo | 1:200 |
|  | CD68 (mouse) | Transmembrane glycoprotein expressed by cells of monocyte lineage, circulating macrophages, and tissue macrophages. | PG-M1 | ab783 | Abcam | 1:200 |

**Supplemental Table II.** Multiplexed staining protocol.

|  | Retrieval | Blocking | Primary Ab | Primary Ab 2 | ImmPRESS | ImmPRESS 2 |
| --- | --- | --- | --- | --- | --- | --- |
| **Hematoxylin** |  |  |  |  |  |  |
| **Cycle 1** | Citra | Horse Serum | Mouse anti-PDPN |  | anti-Mouse |  |
| **Cycle 2** | TRS | Horse Serum | Rabbit anti-LYVE-1 | Rat anti-MECA79 | anti-Rabbit | anti-Rat |
| **Cycle 3** | Citra | Horse Serum | Mouse anti-CD45 |  | anti-Mouse |  |
| **Cycle 4** | Citra | Horse Serum | Rabbit anti-CD20 |  | anti-Rabbit |  |
| **Cycle 5** | Citra | Horse Serum | Mouse anti-CD68 | Rabbit anti-αSMA | anti-Mouse | anti-Rabbit |
| **Cycle 6** | Citra | Horse Serum | Mouse anti-S100 |  | anti-Mouse |  |
| **Cycle 7** | Citra | Horse Serum | Mouse anti-CD8 |  | anti-Mouse |  |
| **Cycle 8** | Citra | Horse Serum | Mouse anti-CD34 | Rabbit anti-AQP1 | anti-Mouse | anti-Rabbit |
| **Cycle 9** | Citra | Horse Serum | Mouse anti-panCK |  | anti-Mouse |  |
| **Hematoxylin** |  |  |  |  |  |  |

**Supplemental Figure 1.**
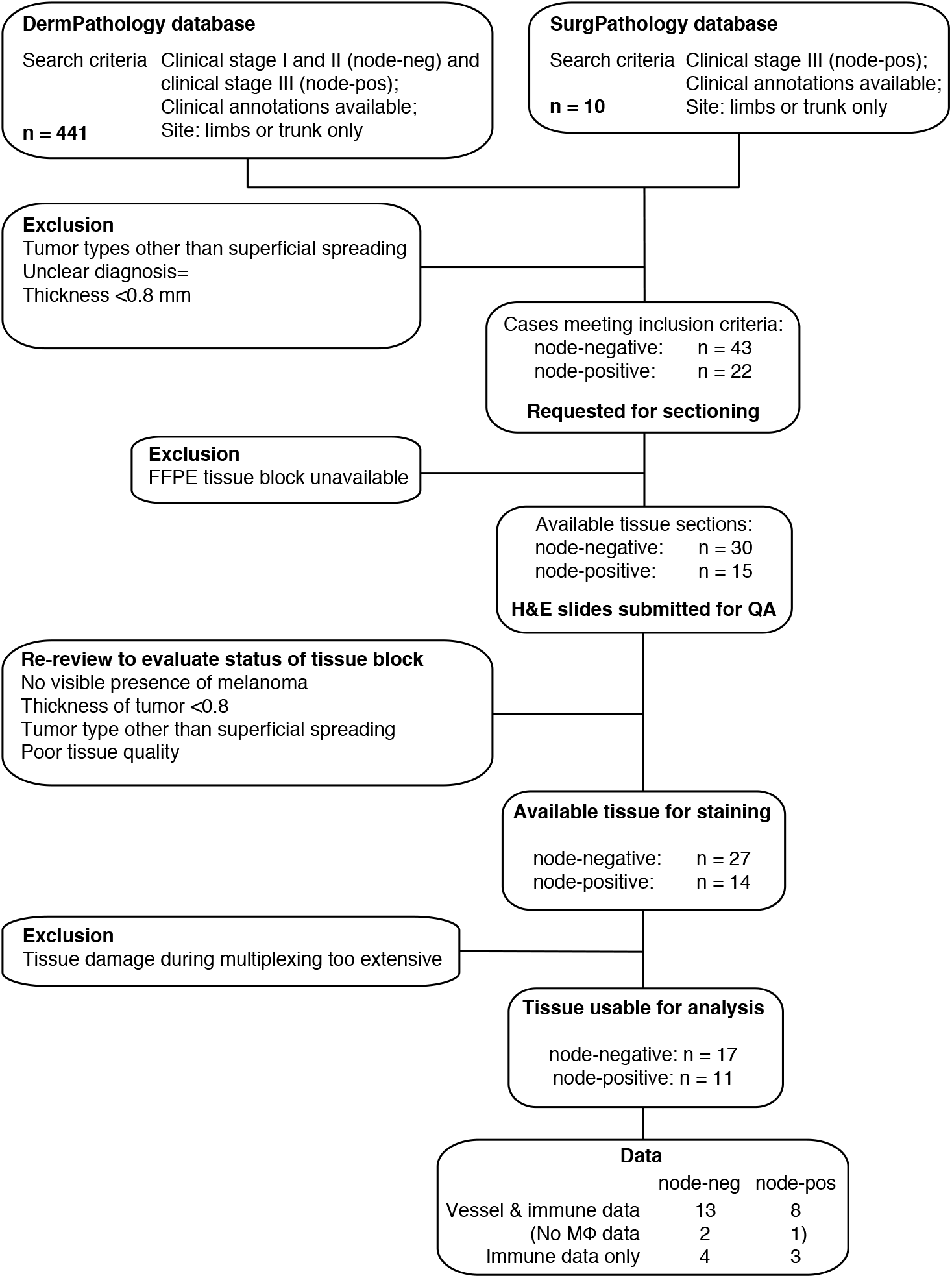
Sample selection and validation flow chart. Sample selection decision tree including criteria for sample inclusion and exclusion. The final cohort consisted of 28 patients. Matched analysis of lymphovasculature and immune infiltrates was performed in 21 patients, with immune data only in an additional 7.

**Supplemental Figure 2.**
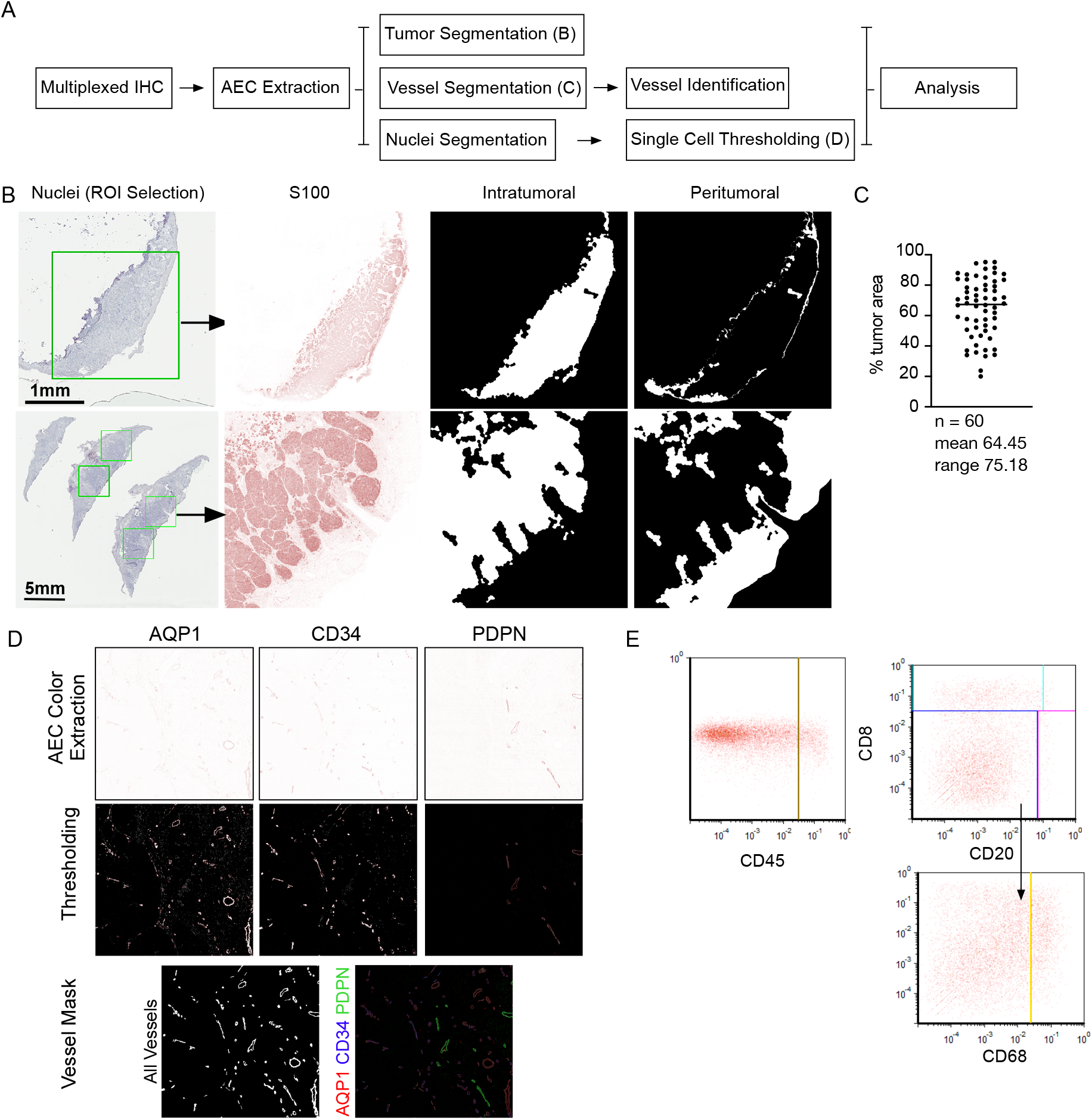
Image analysis workflow. (**A**) Schematic of image analysis workflow. Multiplexed immunohistochemistry (IHC), color (AEC) extraction, image segmentation, single cell and vessel identification and analysis. (**B**) Representative images of nuclei scans and ROI selection, S100 staining and peritumoral and intratumoral segmentation masks. (**C**) Quantitation of percent tumor area per ROI. (**D**) Representative images of color extracted (AEC) single stains for AQP1, CD34, and PDPN, binary thresholding, and generation of vessel masks through combination of all three markers. (**E**) Representative leukocyte gating performed in FCS Express 6 Image Cytometry

**Supplemental Figure 3.**
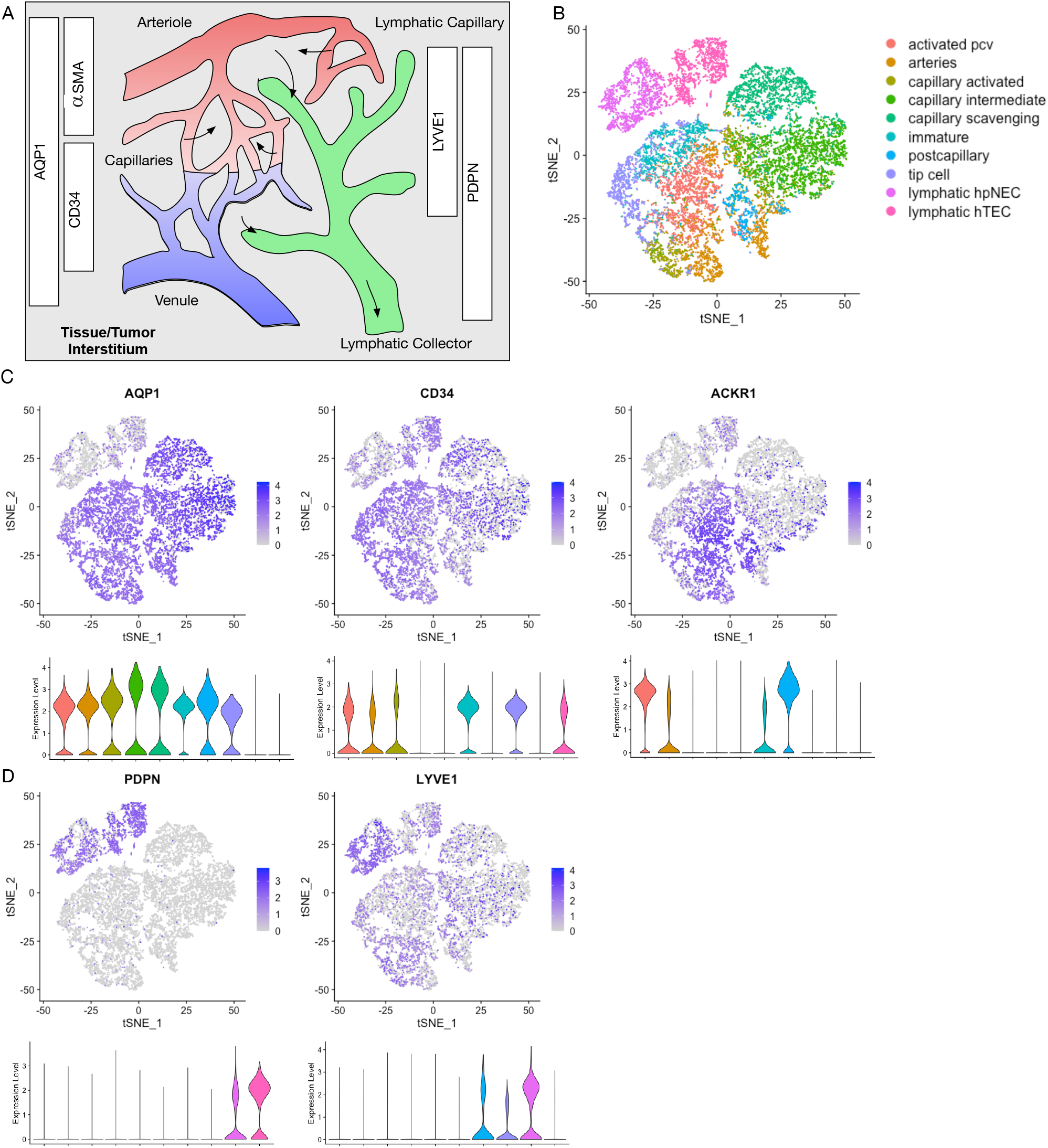
Lymphovascular phenotype specification. (**A**) Schematic representing lymphovascular subtypes identified through imaging workflow. (**B**) Endothelial subtypes identified by single cell sequencing of CD31^+^ cells from lung tumors and adjacent normal tissue (analyzed from Goveia et al *Cancer Cell* 2020). ACKR1 marker of high endothelial venules. pcv= postcapillary venule; hpNEC=human patient normal endothelial cell; hTEC=human tumor endothelial cell. (**C**) Blood and (**D**) lymphatic vessel marker expression across endothelial subtypes.

**Supplemental Figure 4.**
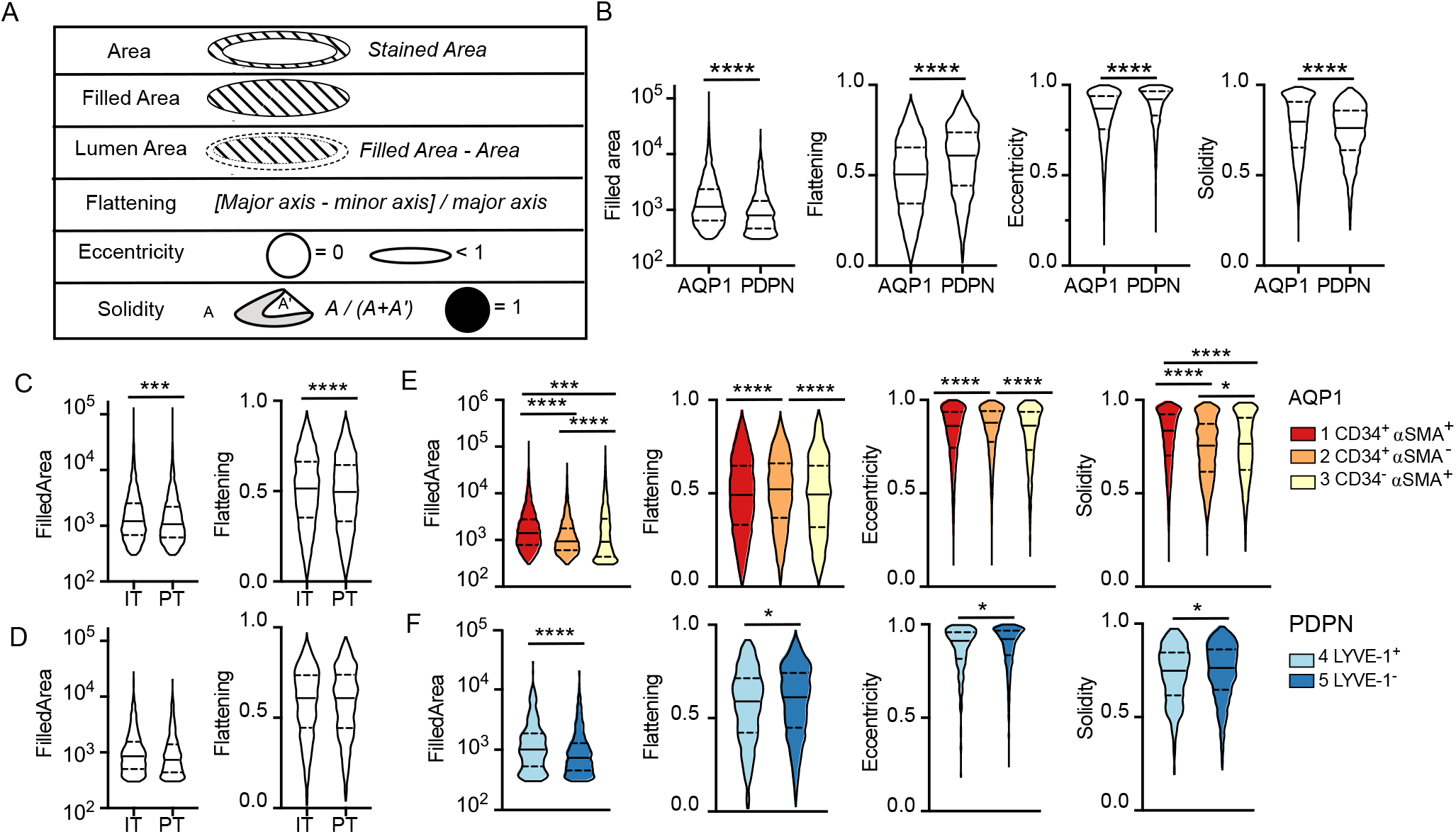
Quantitative morphological features of tumor-associated vessels. (**A**) Schematic explanation of quantitative features extracted from vessel masks. (**B**) All AQP1 or PDPN positive vessel morphological features across all tumor tissue. (**C**) AQP1^+^ vessel morphology in peritumoral (PT) and intratumoral (IT) tumor regions. (**D**) PDPN^+^ vessel morphology in PT and IT tumor regions. (**E**) AQP1^+^ vessel morphology as a function of subtype. (**F**) PDPN^+^ vessel morphology as a function of subtype. Data is representative of individual vessels across samples. Data tested for normality. Mann Whitney test, One-way ANOVA for multiple comparisons. *p<0.05, ***p<0.001, ****p<0.0001.

**Supplemental Figure 5.**
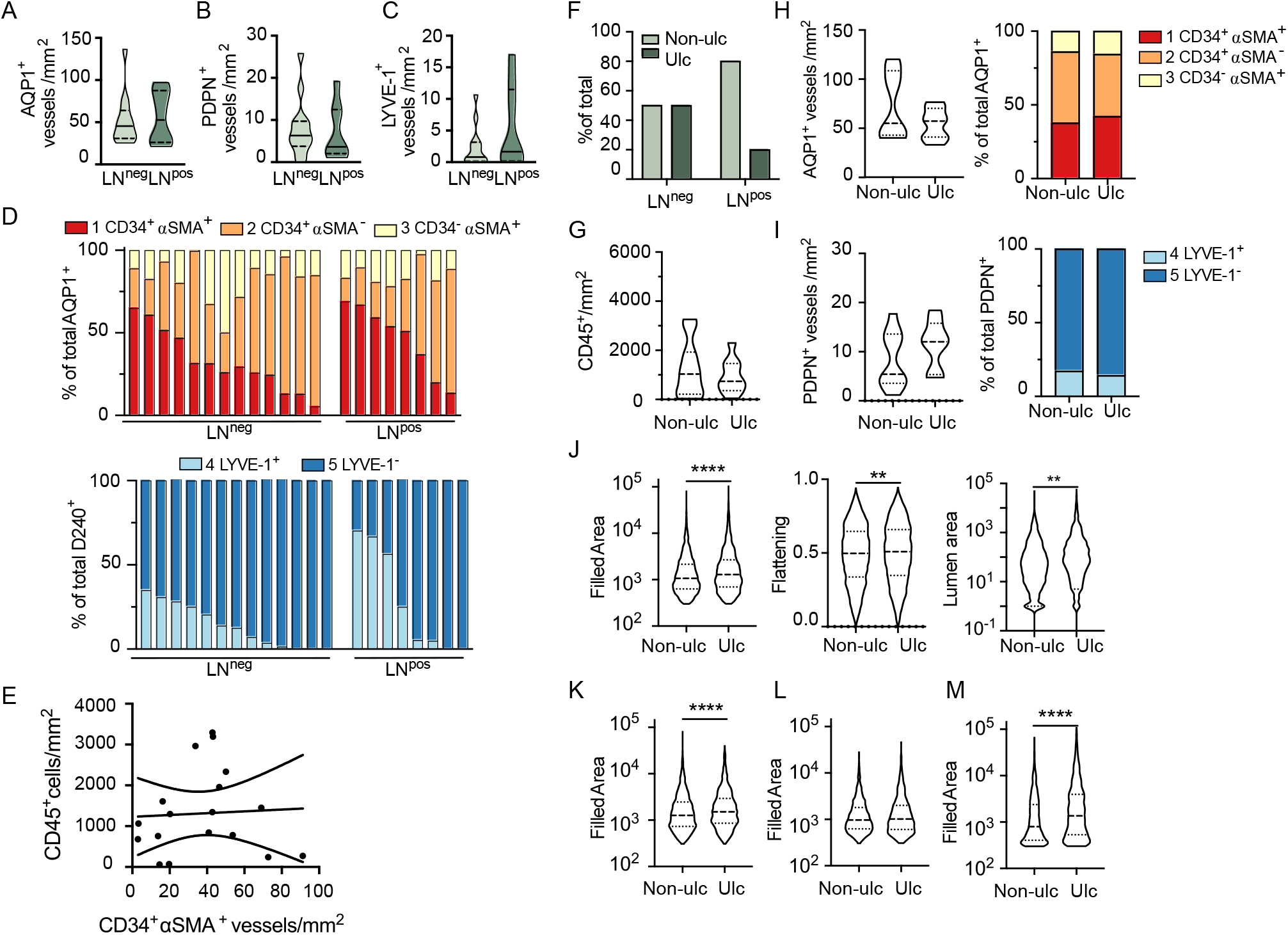
Lymphovascular heterogeneity and ulceration in primary melanoma. (**A**) Intratumoral (IT) AQP1^+^ blood and (**B**) PDPN^+^ lymphatic vessel density. (**C**) Peritumoral LYVE1^+^PDPN^+^ lymphatic vessel density. (**D**) AQP1^+^ blood (top) and PDPN^+^ lymphatic (bottom) vessel heterogeneity per patient in lymph node (LN) positive and negative disease. (**E**) Correlation between CD45^+^ cell density and peritumoral AQP1^+^CD34^+^αSMA^+^ immature capillaries. (**F**) Ulceration (Ulc) as a function of LN status (Non-ulc = non-ulcerated). (**G**) CD45^+^ cell density. (**H**) Total AQP1^+^ blood vessel density and heterogeneity and (**I**) PDPN^+^ lymphatic vessel density and heterogeneity as a function of ulceation. (**J**) Morphological features (filled area, flattening, and lumen area) of all AQP1^+^ blood vessels and filled area in (**K**) CD34^+^αSMA^+^, (**L**) CD34^+^αSMA^-^, and (**M**) CD34^-^αSMA^+^ vessel subtypes. Data is representative of individual vessels. Unpaired student’s t-test, **p<0.01, ****p<0.0001.

**Supplemental Figure 6.**
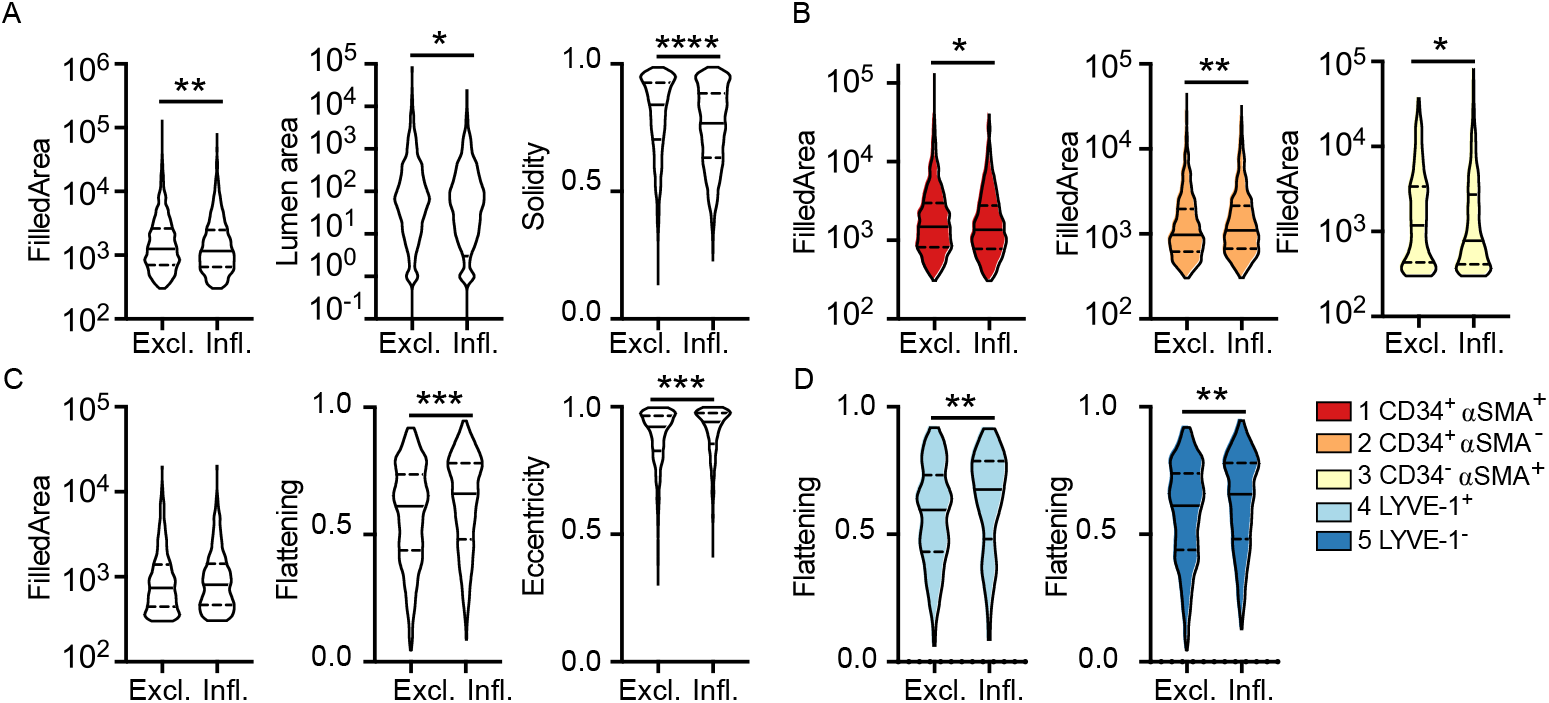
Vessel morphology as a function of CD8^+^ T cell localization in primary melanoma. (**A**) Intratumoral AQP1^+^ vessel features in excluded (Excl.) and infiltrated (Infl.) tumors. (**B**) Vessel size (filled area) as a function of subtype. (**C**) Peritumoral PDPN^+^ vessel features in excluded and infiltrated tumors. (**D**) Vessel flatness as a function of subtype. Data is representative of individual vessels across samples. Data tested for normality. Mann Whitney test, One-way ANOVA for multiple comparisons. *p<0.05, **p<0.01, ***p<0.001, ****p<0.0001.

